# Digital PCR to quantify ChAdOx1 nCoV-19 copies in blood and tissues

**DOI:** 10.1101/2021.05.28.446155

**Authors:** Anita Badbaran, Reiner Mailer, Christine Dahlke, Jannis Woens, Anahita Fathi, Sibylle C. Mellinghoff, Thomas Renné, Marylyn M. Addo, Kristoffer Riecken, Boris Fehse

**Affiliations:** Dept. of Stem Cell Transplantation, University Medical Center Hamburg-Eppendorf (UKE), Hamburg; Institute of Clinical Chemistry and Laboratory Medicine, University Medical Center Hamburg-Eppendorf (UKE), Hamburg; Division of Infectious Diseases, 1st Department of Medicine, University Medical Center Hamburg-Eppendorf (UKE), Hamburg; Department for Clinical Immunology of Infectious Diseases, Bernhard Nocht Institute for Tropical Medicine, Hamburg, Germany; German Center for Infection Research (DZIF), Partner Site Hamburg-Lübeck-Borstel-Riems, Germany; Research Dept. Cell and Gene Therapy, UKE, Hamburg, Germany; University of Cologne, Faculty of Medicine and University Hospital Cologne, Dept. I of Internal Medicine, Excellence Center for Medical Mycology (ECMM), Cologne, Germany; German Centre for Infection Research (DZIF), Partner Site Bonn-Cologne, Cologne, Germany

**Author notes:** Corresponding author:* Prof Dr Boris Fehse, Research Department Cell and Gene Therapy, Department of Stem Cell Transplantation, University Medical Centre Hamburg-Eppendorf, Martinistraße 52, Campus Research (N27), 20246 Hamburg, Germany, Mail.

**Keywords:** SARS-Cov2, ChAdOx1 nCoV-19, vaccination, digital PCR, adenoviral vector

## Abstract

Vaccination with the adenoviral-vector based Astra Zeneca ChAdOx1 nCov-19 vaccine is efficient and safe. However, in rare cases vaccinated individuals developed life-threatening thrombotic complications, including thrombosis in cerebral sinus and splanchnic veins. Monitoring of the applied vector *in vivo* represents an important precondition to study the molecular mechanisms underlying vaccine-driven adverse effects now referred to as vaccine-induced immune thrombotic thrombocytopenia (VITT). We previously have shown that digital PCR is an excellent tool to quantify transgene copies *in vivo*. Here we present a highly sensitive digital PCR for in-situ quantification of ChAdOx1 nCoV-19 copies. Using this method, we quantified vector copies in human serum 24, 72 and 168 hours post vaccination, and in a variety of murine tissues in an experimental vaccination model 30 minutes post injection. We describe a method for high-sensitivity quantitative detection of ChAdOx1 nCoV-19 with possible implications to elucidate the mechanisms of severe ChAdOx1 nCov-19 vaccine complications.

## Introduction

Vaccination has been shown to be effective against severe acute respiratory syndrome coronavirus-2 (SARS-CoV-2).^1^ Adenoviral-vector based vaccines represent one of the cornerstones of the ongoing vaccination programs worldwide. Unfortunately, in contrast to their good safety profile, single cases of severe thromboembolism, often combined with thrombocytopenia were observed for the adenoviral-vector based vaccines from Astra Zeneca (ChAdOx1 nCov-19)^2–4^ and, even more rarely, Johnson and Johnson (Ad26.COV2.S).^5^ By beginning of April 2021, EudraVigilance reported a total of 169 cases of cerebral venous sinus thrombosis (CVST) and 53 cases of splanchnic vein thrombosis after vaccination with ChAdOx1 nCov-19.^6^ This severe complication now referred to as vaccine-induced immune thrombotic thrombocytopenia (VITT, synonym TTS) to some extent resembles atypical heparin-induced thrombocytopenia (HIT) involving platelet-activating antibodies against platelet factor (PF) 4.^3,4^

Vector spread to distinct tissues might contribute to acute, but potentially also long-term side effects, e.g. caused by spontaneous insertion events.^7^ Therefore, it is critical to develop methods that facilitate sensitive analysis of ChAdOx1 nCov-19 vector distribution. A better understanding might help to develop screening approaches for the progression of severe and serious adverse events and thus support prevention and early intervention measures.

We reasoned that digital PCR^8^ should be well suited to detect and quantify vector copies after vaccination in humans, but also in experimental animal models. We previously showed applicability of dPCR to the detection of gene-modified cells in humans.^9^

## Methods

### Decryption of parts of the codon-optimized Spike sequence

To decipher a part of the primary DNA sequence of the Spike coding sequence we designed suitable primers for nested PCR based on the known amino-acid sequence. 200 ng of genomic DNA isolated from blood cells drawn 45 min after vaccination with ChAdOx1 nCov-19 were used as template for the first reaction, 5 μl of a 1:50 dilution for the second. Both first and nested PCRs were performed in 100 μl final volume using Platinum PCR Supermix (ThermoFisher, Kandel, Germany) following a 40-cycle protocol with the following conditions: Initial denaturation: 94°C for 2 minutes; first 10 cycles: 94°C for 30 seconds, 50°C for 45 seconds, 72°C for 30 seconds; next 30 cycles: annealing temperature changed to 60°C; final extension - 7 minutes. Both primary and nested PCR products were visualized on an agarose gel. A single band of the expected size (360 bp) was found after nested PCR and Sanger sequenced (Eurofins, Ebersberg, Germany). The DNA sequence located between the primers and the corresponding amino-acid sequence obtained by *in-silico* translation are depicted in Figure 1. As evident, the amino-acid sequence showed 100% homology to the respective part of the published Spike sequence^10^ (Figure 1).

**Figure 1:**
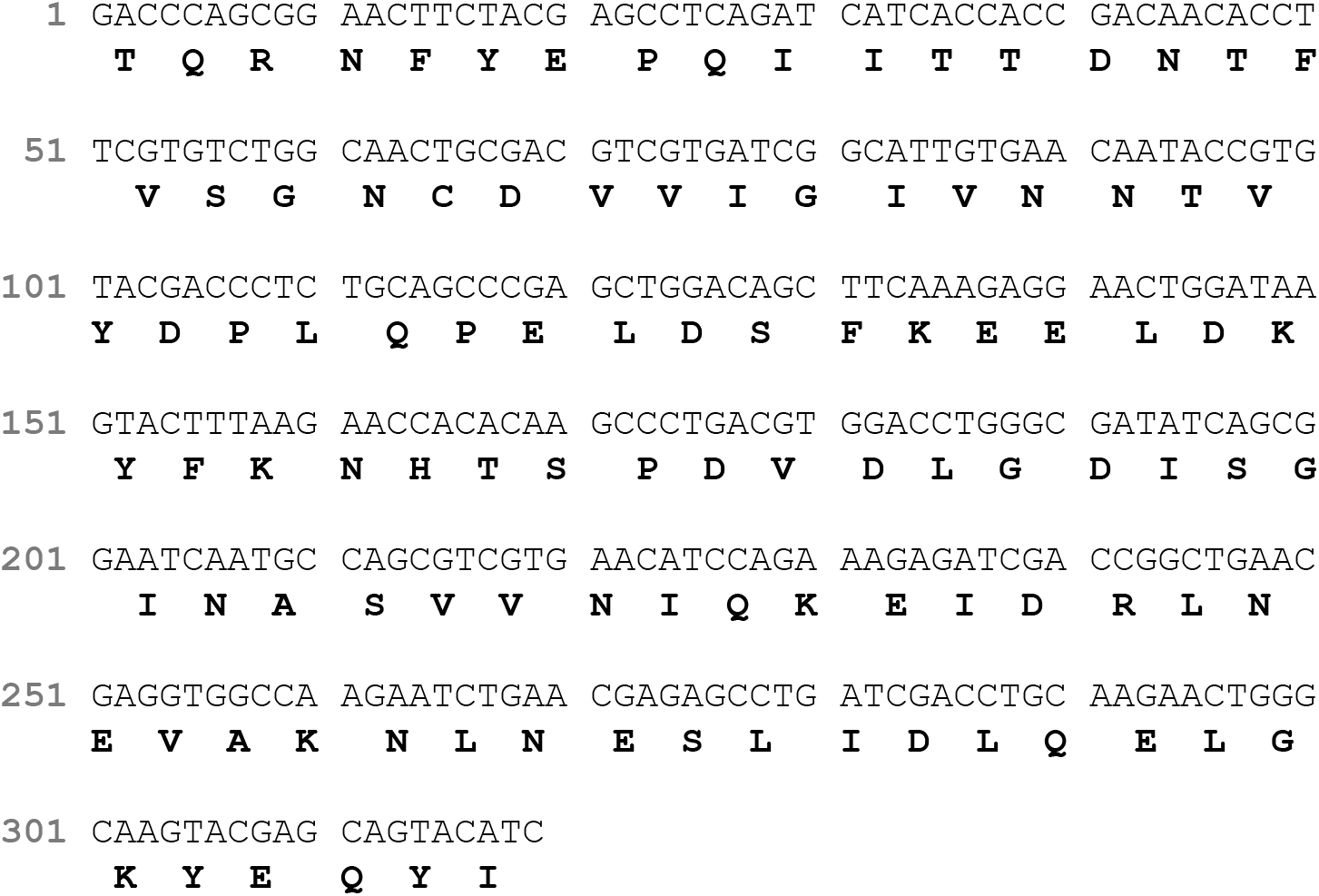
Primary sequence and *in-silico* translation of the identified DNA of the ChAdOx1 nCov-19 vector. The shown amino acid sequence is identical with amino acids 1105 – 1210 of the published Spike protein sequence.^10^

### Design of primers and probe for digital PCR

PCR primers and probes were designed with Primer Express version 3.0.1 (Thermo Fisher). All primers and probes were purchased from Eurofins.

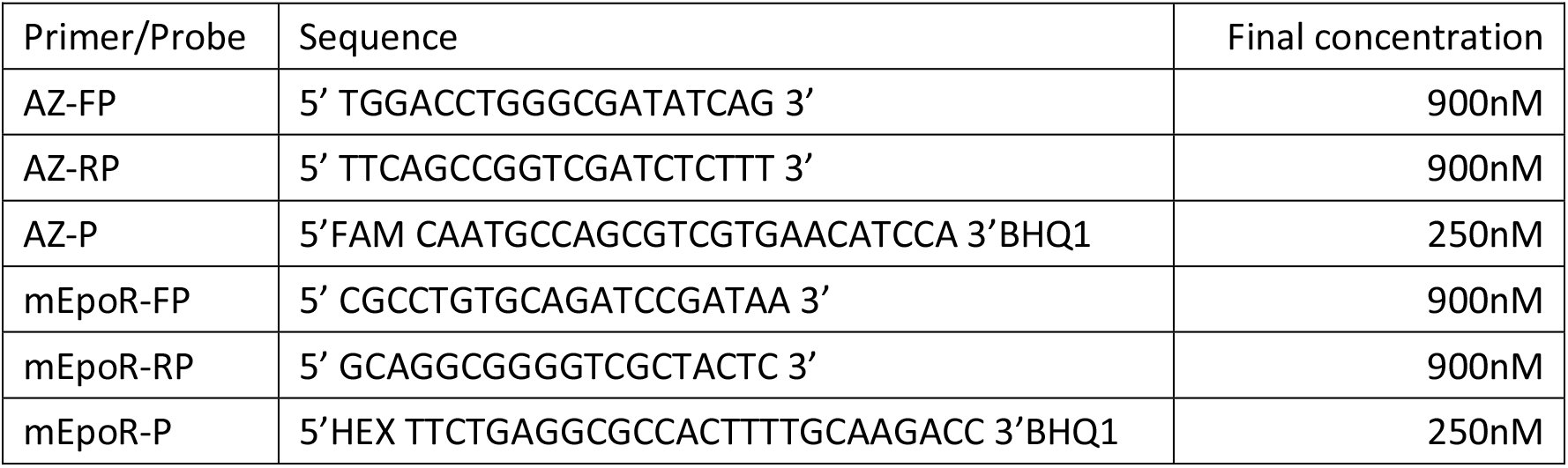

### Isolation of genomic DNA (gDNA) and cell-free DNA (cfDNA)

Genomic DNA (gDNA) from human blood cells was isolated using the QIAamp Blood kit (QIAGEN, Hilden, Germany). To isolate gDNA from different mouse tissues we made use of the blackPREP Rodent Tail DNA Kit (Analytik Jena, Jena, Germany). Cell-free cfDNA (both human and mouse) was isolated with QIAamp DSP Circulation NA Kit (QIAGEN, Hilden, Germany) following the manufacturer’s protocol. 1 to 1.5 ml of human plasma and 0.5 ml murine plasma were used for cfDNA isolation, which was eluted in 50 μl. Measured ChAdOx1 nCov-19 copy numbers were individually corrected to copies per 1 ml plasma.

### Digital-PCR protocol

Digital PCRs were carried out as duplex reactions and analyzed with the QX100 Droplet Digital PCR System (Bio-Rad, California).^9,11^ For human samples, an assay detecting the human ribonuclease P/MRP subunit p30 RPP30 gene (Bio-Rad Assay ID dHsaCP2500350) was used as reference (probe: HEX-BHQ1); for murine cells we included our in-house established reference assay detecting the murine EpoR gene (based on ^12^). In order to investigate gDNA amounts >60 ng, we added 5 U of the restriction enzymes MseI and ApaI (both Thermo Fisher) per individual reaction for human gDNA and mouse gDNA, respectively. PCR conditions followed the general Bio-Rad recommendations.^9^ Data was analyzed with QuantaSoft_v1.7 software (Bio-Rad) including automatic Poisson correction.^9,11^

### Digital-PCR testing

In order to assess sensitivity and specificity of the dPCR we used a dilution series of the PCR product of the first amplification round of the nested PCR; 16 different concentrations were tested. Those samples with expected copy numbers below 10 were tested in triplicates, and mean values were used to build the dilution curve. To test specificity, ten gDNA samples from healthy, non-vaccinated donors were tested in the duplex dPCR - all were highly positive for the REF gene, but negative for ChAdOx1 nCov-19.

### Volunteer samples

Blood samples from nine volunteers were collected at the University Medical Center Hamburg-Eppendorf after their prime vaccination. Donors provided written informed consent (Ethical approval: PV4780). Volunteers received the vaccine ChAdOx1 nCov-19. At days 1, 3 and 7 post vaccination, EDTA-blood was drawn and centrifuged at 200 x *g* for 10 min at RT. Afterwards, plasma was transferred and was again centrifuged at 1000 x *g* for 15 min at RT. Supernatant was stored at −80 °C.

### Mouse model

C57Bl/6 mice of both sexes were anesthetized by intraperitoneal injection of ketamine (120 μg/g BW) and xylazine (16 μg/g BW in saline, 10 μl/g BW) and 50 μl ChAdOx1 nCov-19 vaccine were intradermally injected into the dorsal region. After 30 min mice were sacrificed by cervical dislocation and tissue samples were collected as described.^13^ Citrated blood samples were centrifuged at 1000 x *g* for 10 min at 4°C, and gDNA was separately isolated from centrifuged cells (incl. thrombocytes) and plasma. Data is shown for blood cells. Mice were treated according to national guidelines for animal care at the animal facilities of University Medical Center Hamburg-Eppendorf and approved by local authorities (#56/18). All procedures were conducted in accordance with 3Rs regulations.

## Results and Discussion

In order to identify an appropriate primer-probe combination we first deciphered parts of the primary DNA sequence of the ChAdOx1 nCov-19 vector. We designed a nested PCR located in a suitable region of the Spike-protein coding sequence ^10^ in the vector. After nested PCR on gDNA from blood cells obtained 45 min post vaccination, we obtained a weak, but distinct signal of the expected size visualized by gel electrophoresis (not shown). Sanger sequencing of the purified amplicon followed by *in-silico* translation confirmed complete identity of the amino acid sequence encoded by the PCR fragment with the respective part (amino acids 1105-1210) of the published SARS-CoV-2 Spike protein sequence (Figure 1).^10^

We next designed two PCR primers and an internal dual-labelled hydrolysis probe for dPCR (s. Material and Methods). To verify its sensitivity and specificity, we tested the new assay on negative control samples and a series of dilutions of the first PCR product. We performed duplex-dPCR with a well-established REF gene assay detecting single copies of the human RPP30 gene. As shown in Figure 2a, for both RPP30 and ChAdOx1 nCov-19 vector specific PCRs excellent separation of positive and negative signals was observed. Based on the used dilutions, the highest amount of gDNA tested in our dilution series (approx. 5 ng) corresponds to approx. 750 cells or 1500 haploid genomes. This corresponds very well to the maximal number of RPP30 signals in the dilution series (Figure 2b). As evident, the number of Spike-amplicons (generated in the first PCR round) was approx. 6 times higher in the undiluted sample. Notably, this ratio remained constant for all 16 dilutions tested resulting in a high correlation of the two measured parameters (R2 = 0.9991, p<0.0001; Figure 2c). Moreover, single copies of the targeted Spike-amplicon were readily detected in the assay (Figure 2c), whereas all control samples (gDNA from ten non-vaccinated donors) were negative (not shown). This indicated high sensitivity and specificity of the dPCR assay.

**Figure 2:**
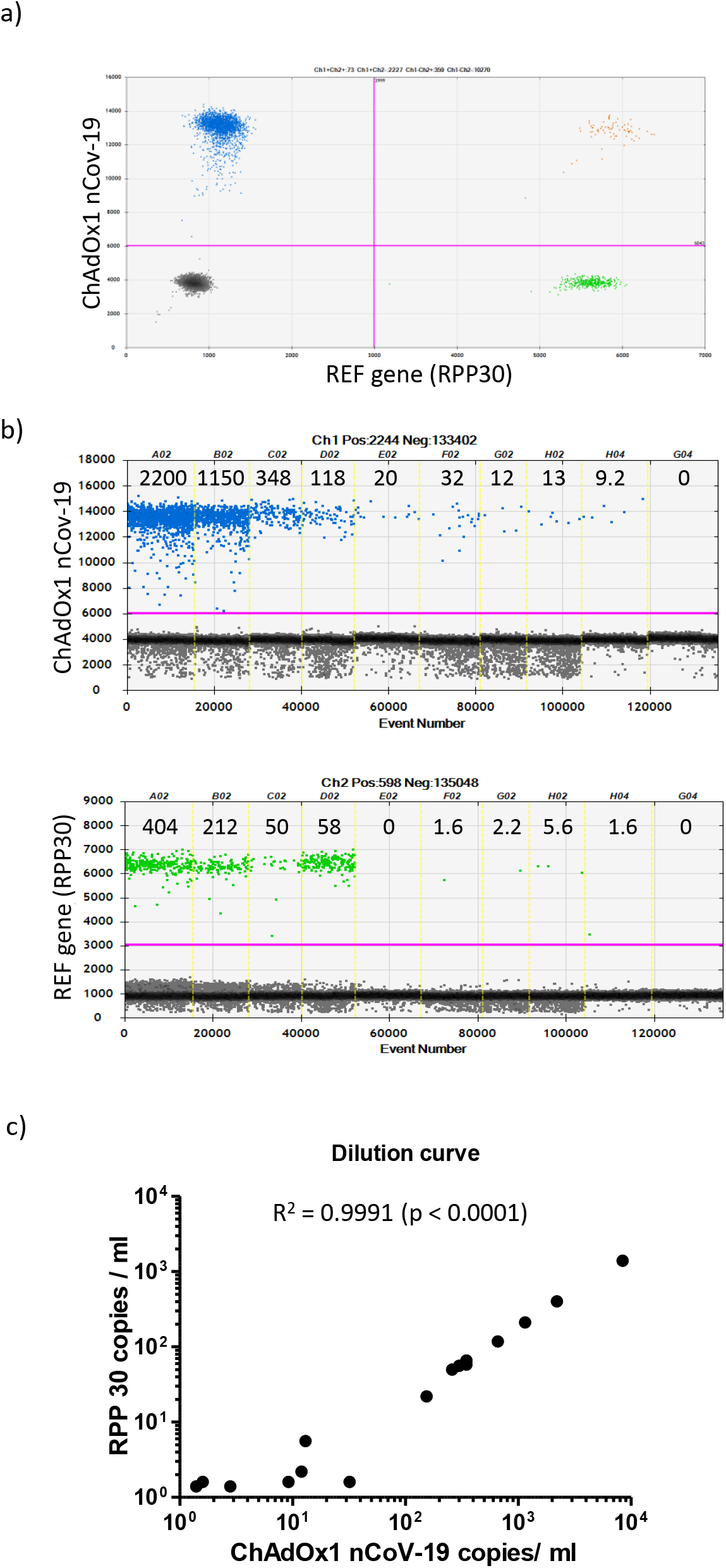
The new dPCR assay facilitates sensitive detection of ChAdOx1 nCov-19 vector DNA. A) The depicted 2D-plot illustrates results of duplex dPCR simultaneously quantifying ChAdOx1 nCov-19 and RPP30 (REF gene) copies. As evident, positive and negative signals are clearly separated. B) Quantification of signals of individual dilutions for the presence of ChAdOx1 nCov-19 (upper panel) and REF (lower panel) copies. Indicated numbers are Poisson corrected. C) Highly significant correlation of ChAdOx1 nCov-19 and REF copy numbers in the dilution series. 16 different dilutions were tested. For dilutions containing less than 10 REF copies, dPCRs were performed in three independent replicates. R^2^ and p values are shown for a two-tailed test.

In order to address practical utility of the assay we tested plasma samples from nine individuals obtained 24, 72 and 168 hours post vaccination. We isolated cell-free cfDNA from blood plasma for each time point. DPCRs were performed in duplex reactions using the RPP30 gene as reference to verify presence of cell-free DNA. At both 24 and 72 hours post vaccination, we detected vector copies in all nine plasma samples, which contained variable amounts of cfDNA (not shown). Interestingly, in six of the nine individuals vector copy numbers were higher 72 hours as compared to 24 hours post vaccination (Figure 3a). Expectedly, no ChAdOx1 nCov-19 vector DNA was found in any of the plasma samples 168 hours post vaccination. All analyses were repeated three times in independent experiments (technical triplicates), mean values are shown in Figure 3a. Taken together, mean copy numbers were more than two times higher at 72 hours as compared to 24 hours post vaccination. In summary, this data proves the usefulness of the presented dPCR assay to monitor the presence of ChAdOx1 nCov-19 vector derived DNA in human blood after regular vaccination.

**Figure 3:**
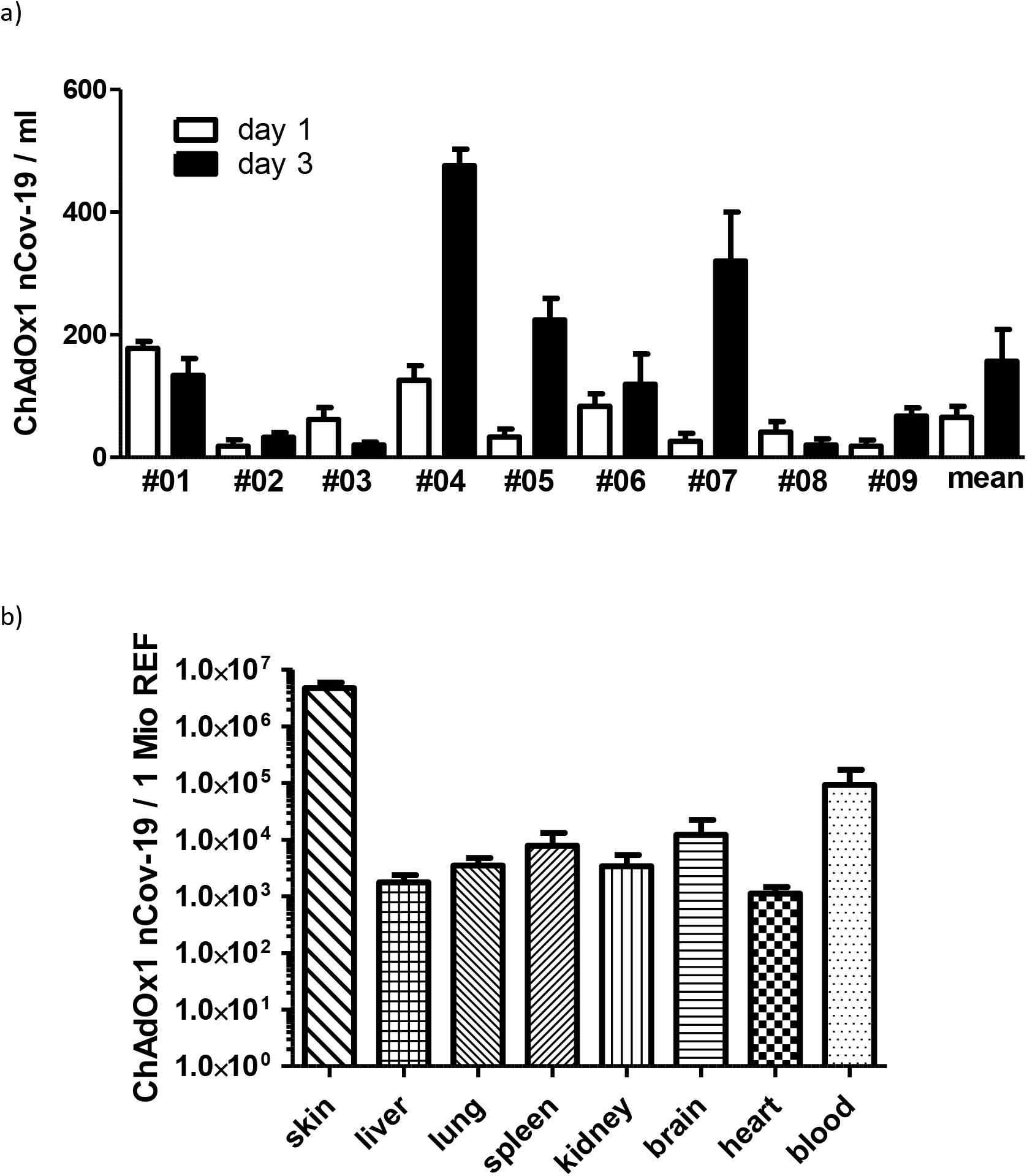
The novel dPCR assay quantifies ChAdOx1 nCov-19 DNA in human post-vaccination plasma samples and murine tissues in an experimental vaccination model. A) ChAdOx1 nCov-19 copies were determined in human plasma from nine volunteers 24, 72 and 168 hours post vaccination. Copy numbers were higher at 72 hours in six out of the nine individuals. No copies were found in the blood 168 hours post vaccination. Mean values and standard deviations (SDs) are shown for triplicate analyses. B) Vector distribution in different mouse tissues 30 minutes after subcutaneous injection using an artificial vaccination model.^13^ Data represents mean values and SDs from four animals. All values for individual animals were measured in independent replicate (2-3) analyses. REF = copies of the reference gene EpoR.

Furthermore, we evaluated whether dPCR might also be applied to detect ChAdOx1 nCov-19 vector copies in different tissues after experimental vaccination. We intradermally injected 50 μl of ChAdOx1 nCov-19 vaccine into the dorsal skin of C57Bl/6 mice.^13^ Challenged animals were sacrificed 30 min later, organs were excised and genomic DNA was isolated from distinct organs/tissues as indicated in Figure 3. Duplex dPCR was performed using the murine *epoR* gene as reference. As expected, huge numbers of ChAdOx1 nCov-19 vector copies were found at the injection site. Detection of ChAdOx1 nCov-19 vector copies in all tissues analyzed provided evidence for imminent vector spread into the blood stream and different organ tissues (Figure 3b).

In summary, we have developed a dPCR assay that facilitates sensitive quantification of ChAdOx1 nCov-19 DNA copies. The assay was successfully applied to quantify ChAdOx1 nCov-19 copies in plasma of vaccinated humans, and also in a variety of tissues in experimentally vaccinated mice. In six of nine individuals we observed a somewhat unexpected kinetics with higher ChAdOx1 nCov-19 copy numbers 72 hours post vaccination as compared to the 24-hour value. This finding might reflect delayed release of ChAdOx1 nCov-19 vectors from the injection site to the bloodstream. Alternatively, cell-free vector DNA in the plasma might in principle also result from the destruction of transduced, Spike-expressing cells by cytotoxic T cells. This question deserves further investigation. In the mouse model, intradermal injection of hyper-physiological doses of ChAdOx1 nCov-19 resulted in rapid distribution of the vector to multiple tissues, including the brain. Determination of the actual (intra- or extracellular) localization of the vector will require future exploration.

The dPCR assay established here might be very useful to study the impact of ChAdOx1 nCov-19 vector distribution *in vivo* with implications for development and kinetics of side effects, both in vaccinated humans and experimental animal models.

## Acknowledgements

We thank Hanna Thode for expert technical assistance and the laboratory team at the UKE, especially My Linh Ly, Monika Friedrich and Niclas Reneviér for collecting and processing the blood from volunteers as well as Amelie Sophie Alberti for study coordination. We acknowledge support by the Deutsche Forschungsgemeinschaft (DFG, German Research Foundation) – grants 125440785 - SFB877 (TR), P6 - KFO306 (TR) and 80750187 - SFB841 (TR & BF) and the German Center for Infection Research (DZIF) - SARS-CoV-2 Fast track fund TTU 01.921 (MMA). Last but not least, the authors also wish to thank all volunteers for their willingness to support our research.

## Author Contributions

BF conceived the study, designed primers and wrote the manuscript, AB isolated DNAs and designed, performed and analyzed dPCRs, RM and TR designed and performed the mouse study, CD, JW, AF, SCM, MMA and KR designed human studies and contributed human samples; all authors reviewed and edited the manuscript.

## Disclosure of Conflicts of Interest

The authors declare no conflict of interest.

## Notes

### Competing Interest Statement

The authors have declared no competing interest.

